# Evolutionary conservation and divergence of regional fructose-responsive gene regulation in the intestine

**DOI:** 10.64898/2026.10.26.761936

**Authors:** Marius Robciuc, Atte Hallasaari, Arto Viitanen, Nicole Lamichane, Ville Hietakangas

## Abstract

Sugar-responsive gene regulation in peripheral organs facilitates safe utilization of fructose. Intestine is the first-line site for fructose catabolism, converting fructose into less harmful metabolites. Its metabolism is spatially organized along the proximodistal axis, but how intestinal zonation shapes responses to fructose and to what extent these responses are evolutionarily conserved remain poorly understood. Here, we examine regional gene regulation in response to acute fructose challenge across the small intestine of mouse and *Drosophila*. Fructose exposure elicited both ubiquitous and regionally biased regulation of metabolic genes. Unlike prolonged feeding on a complex high-carbohydrate diet, fructose feeding activated the heat shock and immune responses, with proximal and distal biases, respectively. The ChREBP/Mondo-Mlx pathway emerged as the major regulator of fructose-responsive gene expression, with a conserved role in regulation of both ubiquitous and region-restricted metabolic targets. Comparative analysis of orthologous genes revealed conserved activation of glucose metabolism, nutrient transport, and stress-responsive pathways, but also notable divergence in downstream metabolic programs. Whereas mouse intestine activates pathways required for fructose conversion into glucose, *Drosophila* preferentially activates de novo lipogenesis. Thus, intestinal gene regulation provides a conserved protective mechanism against high fructose challenge, but the pathways involved have partially diverged during evolution.

## INTRODUCTION

High sugar consumption is a common feature of modern dietary patterns and a major risk factor for metabolic diseases and certain cancers (Adolph and Tilg, 2024). Fructose has attracted particular attention because its excessive consumption is linked to cardiometabolic diseases associated with liver fat accumulation, as well as cancer (Febbraio and Karin, 2021). The intestine plays a critical role in limiting systemic exposure to fructose by metabolizing a substantial fraction of ingested fructose into glucose and organic acids within enterocytes before it reaches the portal circulation (Bismut et al., 1993; Jang et al., 2018). This intestinal “shielding” capacity is effective when fructose is consumed at moderate amounts and gradually, but becomes insufficient following acute high-dose exposure (Jang et al., 2020; Zhao et al., 2020). Studies measuring portal blood fructose concentrations in humans and other mammals indicate that, at high intake levels, a substantial fraction of fructose escapes intestinal clearance and reaches the liver (Herman and Birnbaum, 2021). In mice fed high-doses of fructose, intestinal microbiota converts fructose-derived substrates into acetate that can subsequently contribute to hepatic de novo lipogenesis (Jang et al., 2018; Zhao et al., 2020). Thus, the intestine represents an important metabolic barrier that determines the extent and nature of systemic exposure to dietary fructose.

Intracellular sugar sensing is a conserved mechanism that enables animal cells to adjust gene expression according to nutrient availability. The ChREBP/Mondo-Mlx complex is a heterodimeric basic helix-loop-helix transcription factor that functions as a central regulator of sugar-responsive gene expression in metabolic tissues, including the liver, intestine, and adipose tissue (Iizuka et al., 2004; Li et al., 2006; Havula and Hietakangas, 2012; Herman et al., 2012; Havula et al., 2013; Kim et al., 2017; Cadena Del Castillo et al., 2025). Both its molecular architecture and many of its target genes are conserved across metazoans, from mammals to *Drosophila* (Li et al., 2006; Havula and Hietakangas, 2018). ChREBP/Mondo-Mlx controls metabolic genes in a tissue-specific manner, both directly and through a network of second-tier transcription factors (Havula et al., 2013; Bartok et al., 2015; Mattila et al., 2015; Kokki et al., 2021; van den Berg et al., 2023). Genetic disruption of *Mlxipl*, which encodes ChREBP, in mice or *bigmax*, which encodes Mlx, in *Drosophila* causes severe sugar intolerance and, can result in lethality under high-sugar conditions (Havula et al., 2013; Iizuka et al., 2004). The importance of ChREBP in intestinal fructose handling is further supported by tissue-specific deletion studies showing that ChREBP activity in the small intestine is required for adaptation to high-fructose feeding (Kim et al., 2017), consistent with the role of the intestine as a first-line metabolic buffer for dietary fructose (Jang et al., 2020).

Most intestinal nutrient absorption and metabolism occur in the small intestine, which in mammals is traditionally divided into three anatomical segments: the duodenum, jejunum, and ileum. These segments lack sharply defined anatomical boundaries, and many genes display gradual expression gradients along the intestinal proximal-distal axis (Bates et al., 2002; San Roman and Shivdasani, 2011). Recent analyses of enterocyte transcriptomic profiles have provided a more refined view of this regional organization, identifying five distinct zones of metabolic gene expression along the mouse and human small intestine (Zwick et al., 2024). Importantly, these metabolic domains are nutrient-responsive: one week of feeding a high-carbohydrate diet was sufficient to alter their gene expression, demonstrating substantial transcriptional plasticity in response to dietary changes (Zwick et al., 2024). However, how different intestinal regions respond transcriptionally to acute fructose exposure, and to what extent such regional responses are conserved across species, remain unclear.

Here, we address how regional gene expression along the proximodistal axis of the intestine responds to acute fructose feeding and to what extent these responses are conserved across multicellular animals. We find that acute fructose exposure engages a core transcriptional program conserved between mouse and *Drosophila* that includes metabolic adaptations and distinct stress responses. ChREBP/Mondo-Mlx is a central mediator of the fructose response, controlling both broadly distributed and regionally restricted metabolic genes. Whereas many metabolic responses are activated throughout the intestine, stress responses are spatially segregated, displaying either proximal or distal bias. Despite conservation of responses related to glucose handling, nutrient transport, and cellular stress between mouse and *Drosophila*, gene regulation related to fructose shielding differs between species: in the mouse intestine, fructose strongly activates a transcriptional program involved in fructose conversion to glucose, whereas the *Drosophila* midgut responds through activation of genes associated with *de novo* lipogenesis. In sum, our findings provide a global view of intestinal gene regulation in response to fructose feeding, including its regional organization and evolutionary conservation, and provide insight into the physiological adaptation of the intestine to the metabolic challenge of high fructose exposure.

## RESULTS

### Fructose feeding activates ubiquitous and regional metabolic gene expression in the mouse small intestine

To uncover intestinal gene regulation in response to acute fructose feeding we challenged fasted male mice with one dose of 4g/kg fructose for two hours, applying oral gavage protocols used in previous studies (Kim et al., 2017; Goncalves et al., 2019; Taylor et al., 2021). The challenge was well tolerated, although strong intestinal bloating was observed (**Fig. S1A**). To identify possible regionality of fructose responses, we divided the small intestines into five segments (**Fig. 1A**), following the findings of Zwick and coworkers on zonal metabolic gene expression (Zwick et al., 2024). Villi were harvested by scraping the luminal side of the small intestine, which allowed rapid sample processing for RNA sequencing with an enrichment in enterocyte composition (**Fig. 1A**).

**Figure 1.**
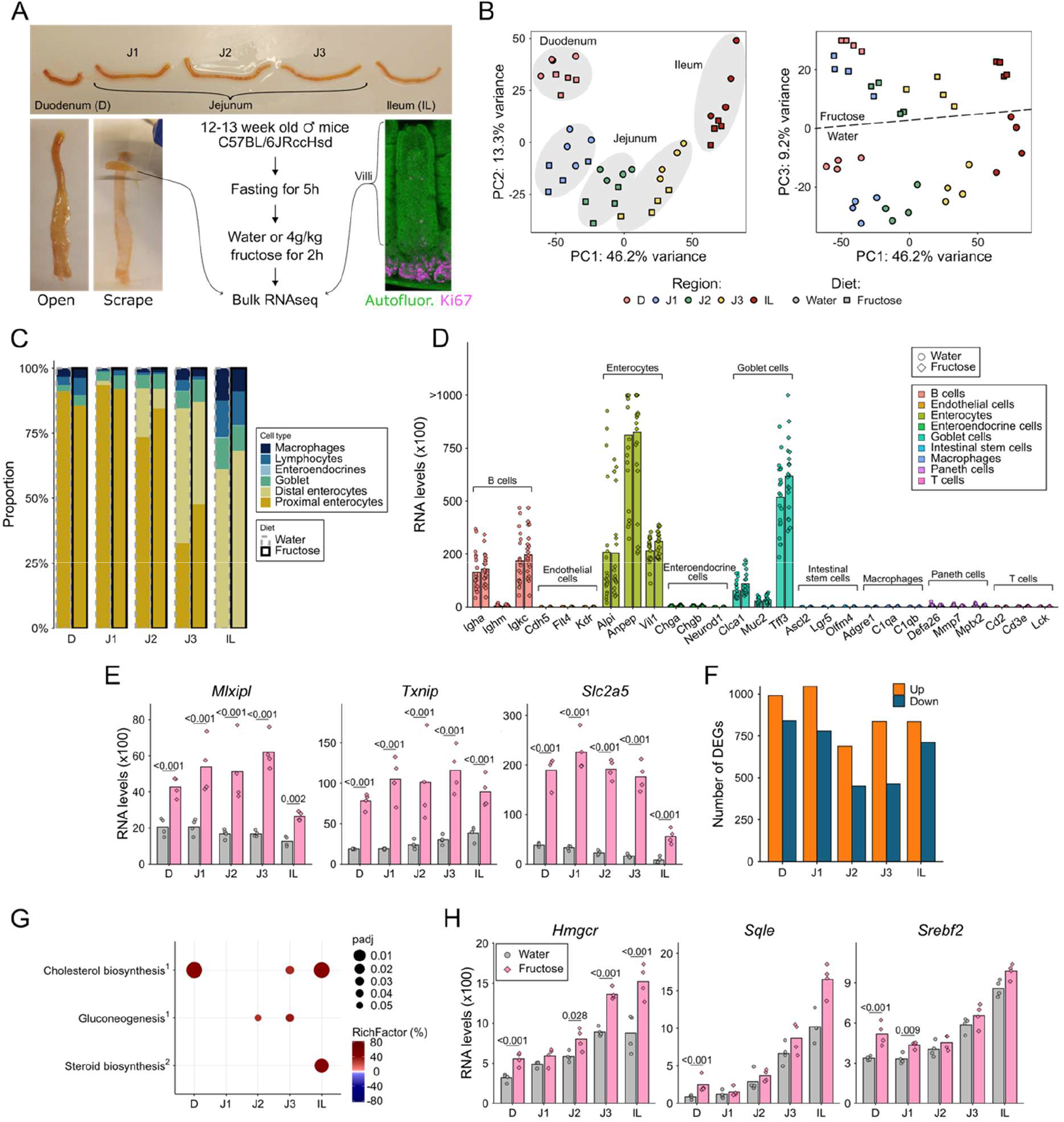
High fructose feeding causes ubiquitous activation of ChREBP-Mlx target genes and regional changes metabolic gene expression in the mouse small intestine. **A)** Sample annotation and preparation, mouse strain details, and experimental design. N = 4 mice per group, used in all downstream analysis. Immunofluorescence image shows the villi region used for bulk RNA-seq and the proliferative crypt area that was excluded from sampling. **B)** Principal component analysis (PCA) showed a clear separation of samples based on harvesting region with 46.2% (PC1) and 13.3% (PC2) of total variance. Samples segregated based on the indicated treatment condition with 9.2% (PC3) of total variance. **C)** Relative cell composition estimates based on deconvolution of bulk RNA-seq data using MuSiC, highlighting the enterocytes as the dominant cell type in the villi and an increased presence of immune cells in the ileum. **D)** Expression levels of marker genes for cell types commonly found in the small intestine. **E)** Expression levels of established ChREBP-Mlx target genes induced by fructose in all regions of the intestine. **F)** Total number of differentially expressed genes (DEGs) upon fructose challenge in each region. **G)** Metabolism related region-specific enriched pathways resulted from over representation analysis (ORA), with padj < 0.05 (dot size). Pathway Richfactors resulting from upregulated DEGs are on a red scale and those resulting from downregulated DEGs were converted into negative values and shown on a blue scale. Databases used were: ^1^Reactome, ^2^KEGG. **H)** Expression levels of selected genes related to cholesterol biosynthesis pathway. For **E** and **H**, p values were determined with DESeq2 and adjusted using Benjamini-Hochberg correction. Adjusted p-values > 0.05 are not shown. N = 4 for all groups.

Principal component analysis (PCA) of the bulk RNA-seq data revealed sample clustering according to region with 46.4% (PC1) and 13.3% (PC2) of total variance (**Fig. 1B**). While all five regions could be separated, the three regions mapping to jejunum clustered closer to each other. Following the conventional anatomical division of the small intestine, we annotated the samples as duodenum (D), jejunum (subdivided into three regions: J1, J2, and J3), and ileum (IL). PCA also revealed clustering according to treatment condition, accounting for 9.2% (PC3) of total variance (**Fig. 1B**). These results show that both regional identity and the response to fructose can be clearly identified based on global gene expression.

We opted to use bulk RNA sequencing of villi in order to achieve a global view to fructose-induced responses, but also considered technical aspects, such as high sequencing depth and minimal sample processing. To estimate the proportion of cell types in our samples, we utilized the existing scRNA-seq data (Chen et al., 2025) and the deconvolution package MuSiC (Wang et al., 2019). Consistent with the expected cell distribution in villi, enterocytes were predominantly represented, accompanied by goblet and immune cells (**Fig. 1C**). Moreover, proximal and distal enterocytes were identified in their respective sampling regions and the proportion of immune cells increased in ileum. Plotting the absolute expression levels of individual marker genes highlight only the genes with very high expression levels in enterocytes, goblet cells and B cells and but not in intestinal stem cells or Paneth cells, located in the crypts (**Fig. 1D**). These data are consistent with the conclusion that our sampling captures the global gene expression of cells present in the villi.

To assess ChREBP activity along the mouse small intestine we selected genes that have previously been shown to respond strongly upon high fructose feeding in a ChREBP dependent manner (Kim et al., 2017). *Mlxipl* (ChREBP encoding gene, autoregulation), *Txnip and Slc2a5* (GLUT5, fructose transporter) genes were all upregulated throughout the intestine albeit somewhat less in the ileum (**Fig. 1E**). The observed lower response in ileum could be due to lower uptake of fructose, suggested by lower basal mRNA levels of fructose transporter GLUT5 (**Fig. 1F**) and ketohexokinase (KHK) (**Fig. S1B**), the enzyme responsible for the intracellular phosphorylation of fructose.

Next, we performed a global unbiased analysis of all differentially expressed genes (DEGs, log2 fold change > 0.5, adj.p < 0.05) and their distribution into biological pathways. In all regions, several hundreds of both upregulated and downregulated DEGs were observed (**Fig. 1F**). Many of the DEGs were unique to each region, with highest numbers in duodenum and ileum and lowest in the middle J2 region (**Fig. S1C**). Region D and J1 had the most shared DEGs when compared with other regions that were close or most distant to each other (**Fig. S3C**), which can be explained by the lack of clearly defined borders between the regions (Zwick et al., 2024). While the number of up- and downregulated DEGs were relatively similar, genes that were shared DEGs between all regions (like the ChREBP targets in Fig. 1F), were predominantly upregulated by fructose feeding (**Fig. S1C**). Over-representation analysis revealed a much higher number of significantly enriched pathways (adj.p < 0.05) resulting from the upregulated DEGs compared to those from downregulated DEGs (**Fig. S1D**). Most of the upregulated pathways were shared between the regions except for IL, which had the most unique pathway composition (**Fig. S1E**).

Surprisingly we found a relatively small number of metabolic pathways that were enriched in our samples. Gluconeogenesis pathway was significantly upregulated in the middle part of the intestine upon high fructose ingestion (**Fig. 1G**). This overlaps with the location where genes related to basal carbohydrate metabolism were previously shown to be enriched (Zwick et al., 2024). The only other significantly enriched metabolism-related pathways were cholesterol and steroid biosynthesis, upregulated at the proximal and/or distal end of the small intestine (**Fig. 1G**). Screening of individual genes involved in cholesterol metabolism revealed *Hmgcr* and *Sqle* increased expression after high fructose feeding and a clear increase in basal expression levels in ileum (**Fig. 1H**). The proximal-distal expression pattern was also observed for SREBP2 transcription factor encoding gene (*Srebf2*) which is a master regulator of *de novo* cholesterol biosynthesis. Strikingly, these genes have the opposite response in liver from mice fed a high fructose diet where these genes were reduced after high fructose diet (Zhang et al., 2017).

Collectively these results show that acute intake of fructose induces an intestine-wide activation of known ChREBP target genes, accompanied by other region-specific transcriptional responses. Although canonical carbohydrate and lipid metabolic pathways were only modestly enriched, the induction of cholesterol biosynthesis genes, particularly in regions with higher basal *Srebf2* expression, revealed an unexpected metabolic response to acute fructose exposure in the small intestine.

### High fructose induces regional stress and immune responses

Next we compared our bulk RNA-seq data with single cell RNA sequencing (scRNA-seq) data from enterocytes in the study by Zwick et al., where mice were chronically (1 week) fed a high carbohydrate diet (Zwick et al., 2024). This comparison allowed us to distinguish responses occurring during prolonged feeding on complex high carbohydrate diet (38% starch, 13% maltodextrin and 17% sucrose) vs an acute fasting to high fructose response (Kim et al., 2017). For this analysis, we converted the scRNA-seq data into pseudo-bulk profiles. High fructose dataset had 22.6% more genes compared to the high carbohydrate diet dataset (**Fig. 2A**). Despite substantial overlap in detected genes, the number of DEGs was markedly higher in the fructose-fed animals, and only 0.9% DEGs shared between the two datasets (**Fig. 2B**). Bearing in mind the limitation that the cell composition of the samples was not identical, the extent of observed differences likely reflects that acute high fructose challenge of fasted animals activates a higher number and distinct set of genes compared to long term feeding of diet with high carbohydrate content.

**Figure 2.**
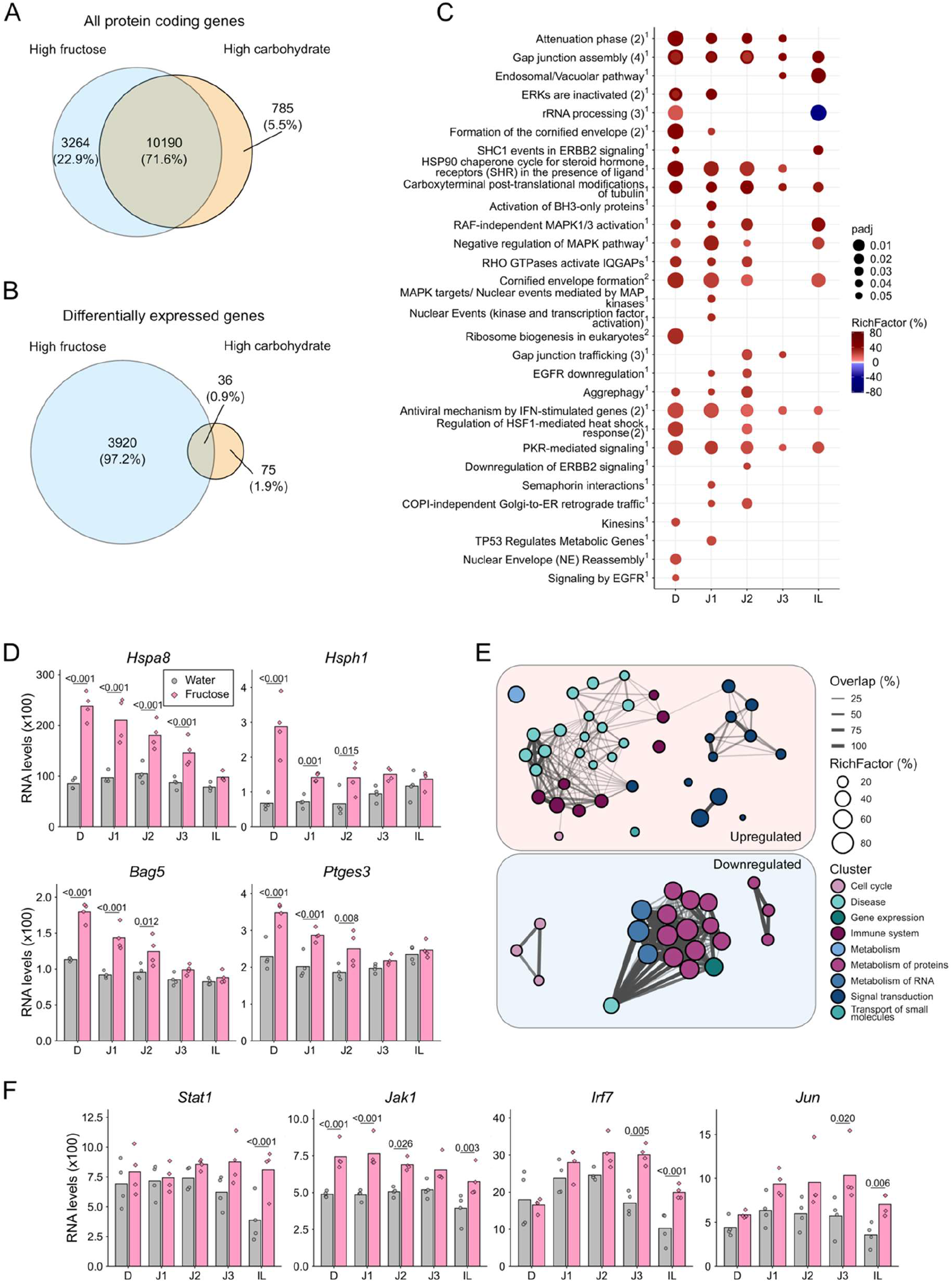
Acute high fructose feeding triggers stress- and immune-related gene responses in a region-specific manner. **A**) Venn diagram with the total number of protein coding genes detected in the high fructose and high carbohydrate datasets. **B**) Venn diagram with the total number of differentially expressed genes in the high fructose and high carbohydrate datasets. **C**) Top 30 regionally enriched pathways by Richfactor resulted from over representation analysis (ORA), with *P* < 0.05 (dot size), adjusted for multiple comparison using the Benjamini–Hochberg correction. Pathway analysis by Richfactor resulting from upregulated DEGs are on a red scale and those resulting from downregulated DEGs were converted into negative values and shown on a blue scale. Databases used were: ^1^Reactome, ^2^KEGG. **D**) Examples of genes related to heat shock response. **E**) All ileum-specific enriched pathways resulted from over representation analysis (ORA), with *P* < 0.05, adjusted for multiple comparison using the Benjamini–Hochberg correction. Each node represents an individual pathway, with node size representing Richfactor. Node colour represents the cluster to which the pathway is annotated to. Width of connecting lines represent the overlap of the sets of differentially expressed genes annotated to the pathways. **F**) Examples of genes related to stress and inflammation. For **D** and **F**, p values were determined with DESeq2 and adjusted using Benjamini-Hochberg correction. Adjusted p-values > 0.05 are not shown. N = 4 for all groups.

Over-representation analysis of the high carbohydrate dataset revealed several metabolism-related pathways in the proximal intestine (**Fig. S2A**). In contrast, the acute high fructose feeding resulted in an enrichment of pathways associated with the heat shock response controlled by the protein-damaging stress-activated transcription factor HSF1 (**Fig. 2C**). Genes regulated by the heat shock response include the Hsp70-family molecular chaperone Hspa8, nucleotide exchange factor Hsph1, Hsp70 co-chaperone Bag5 as well as Ptges3, encoding an Hsp90 co-chaperone p23 (Pearl and Prodromou, 2006; Pustovaya et al., 2026). All these genes were significantly upregulated following the high-fructose challenge, with the strongest increase observed in the duodenum and progressively weaker responses towards the distal small intestine (**Fig. 2D**). Notably, none of these genes were significantly affected by the chronic high carbohydrate diet (**Fig. S2B**).

Ileum displayed the largest number of uniquely altered pathways upon acute high fructose feeding (**Fig. S1E**). Therefore, we examined next whether these pathways converged on common biological processes. Indeed, most of the upregulated pathways in the ileum were associated with innate immune system or diseases characterized by strong immune and inflammatory responses (**Fig. 2E**). Such responses are frequently accompanied by activation of the integrated stress response and suppression of protein synthesis, which can protect cells from further stress (Cláudio et al., 2013; Costa-Mattioli and Walter, 2020). Consistent with this interpretation, most of the downregulated pathways in the ileum were related to RNA and protein metabolism (**Fig. 2E**). Genes encoding the JAK/STAT pathway, a major regulator of inflammatory and stress-responsive gene expression, had also increased expression in the ileum following high-fructose exposure (**Fig. 2F**). Similar increase was observed for Irf7, a key regulator of type I interferon responses, and Jun, transcriptional regulator of stress responses, and pro-inflammatory cytokine expression. None of these genes were significantly affected by the high carbohydrate diet feeding (**Fig. S2C**).

Collectively, these results indicate that a high fructose challenge, but not chronic feeding on complex carbohydrate-rich diet, elicits spatially distinct stress responses, including a heat shock response in the duodenum and an inflammatory response in the ileum.

### Mondo-Mlx is a predominant mediator of metabolic gene expression response to fructose in *Drosophila* larvae

Our previous work has established the *Drosophila* larva as a powerful genetic model to study the sugar-responsive gene regulation by the ChREBP ortholog Mondo and its obligate binding partner Mlx (Havula et al., 2013; Havula and Hietakangas, 2018; Cadena Del Castillo et al., 2025). Staining of the larval midgut (small intestine) with an in-house generated anti-Mlx antibody revealed its nuclear translocation in multiple regions upon high fructose feeding for six hours, consistent with Mondo-Mlx activation (**Figure 3A**). To determine the role of Mondo-Mlx in fructose response along the larval intestine we used *mlx* null mutant animals (*mlx^1^*) and their corresponding controls fed a chemically defined diet without sugar or with 15% fructose for six hours. Pupariation assay showed that *mlx^1^*larvae are highly sensitive to fructose added to holidic media, as even 3% added fructose was sufficient to cause a delay in pupariation for *mlx^1^*larvae compared to control (**Fig. S3A** and **S3B**).

**Figure 3.**
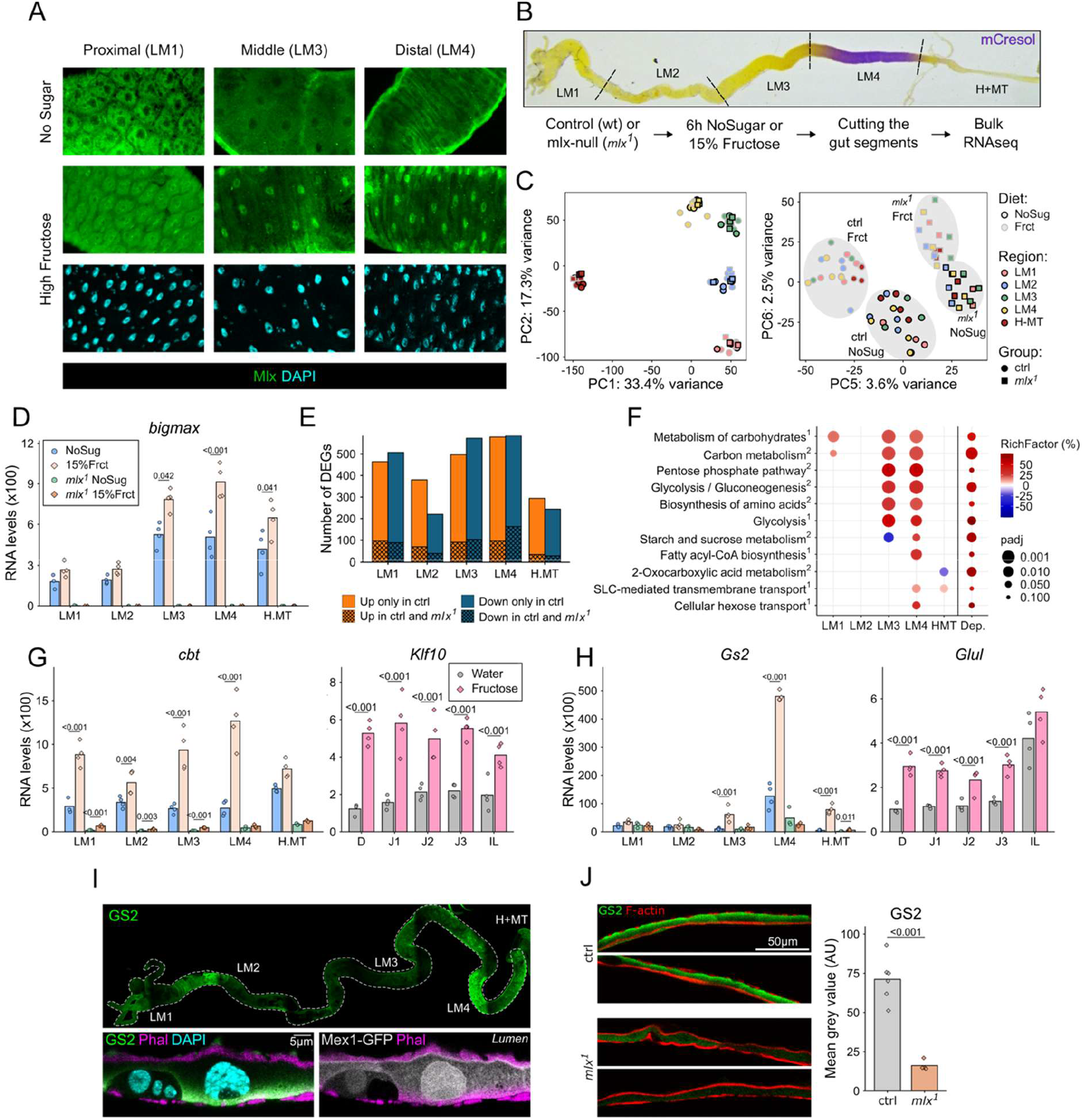
The ubiquitous intestinal response to fructose is conserved in *Drosophila* and mostly dependent on Mondo-Mlx. **A**) Mlx (green) and DAPI (cyan) staining in the indicated regions of the midgut from *Drosophila* larvae fed for 6 hours a holidic diet without sugar or 15% fructose. **B**) Intestine cutting strategy, sample annotation, and experimental design. Groups had 3-4 biological replicates with each replicate containing gut fragments from five larvae. Alkaline region was visualized by addition of mCresol in the experimental diets. **C**) Principal component analysis (PCA) showed a clear separation of samples based on harvesting region with 33.4% (PC1) and 17.3% (PC2) of total variance. Samples segregated into the four experimental groups with 3.6% (PC5) and 2.5% (PC6) of total variance. **D**) Expression levels of the Mlx-encoding gene *bigmax*. **E**) Total number of differentially expressed genes (DEGs) in each region in control genotype. Cross-hatched part of the bars represents the number of DEGs that response similarly in Control and *mlx*-null larvae (*mlx^1^*) fed high fructose. **F**) All regionally enriched Mlx-dependent pathways resulted from over representation analysis (ORA), with *P* < 0.05 and *P* < 0.10 for dependency (dot size), adjusted for multiple comparison using the Benjamini–Hochberg correction. Pathway Richfactor resulting from upregulated DEGs are on a red scale and those resulting from downregulated DEGs were converted into negative values and shown on a blue scale. Databases used were: ^1^Reactome, ^2^KEGG. **G, H)** Examples of orthologous genes that respond similarly to high fructose feeding in *Drosophila* larvae and mouse, in a Mlx-dependent manner. **I**) Colocalisation of GS2 (green) protein expression with enterocyte marker Mex1-GFP (grey) from larvae fed high fructose for 24 hours. Full gut image (top) show GS2 expression in LM4 region and high magnification closeup images (below, scale 5 µm) show the enterocyte expression. F-actin staining (Phalloidin, magenta) marks the gut muscle layer and the enterocyte brush border. **J**) Quantification of GS2 protein expression in LM4 region from Control and *mlx*-null larvae fed high fructose diet for 24 hours and representative images of the guts used in the analysis. P-value calculated using two-tailed Welch’s t-test. N = 4-6, each dot represents a biological replicate. Adjusted p-values > 0.05 are not shown.

Intestinal segmentation was facilitated by the addition of pH sensor mCresol in the larvae food (Overend et al., 2016), which revealed the alkaline region of the distal midgut (**Fig. 3B**). We collected four larval midgut regions (LM 1-4) and a fifth sample combining the hindgut plus Malpighian tubules (H+MT) for bulk RNA-seq analyses (**Fig. 3B**). PCA revealed a clear sample clustering according to the region with 33.4% (PC1) and 17.3% (PC2) of total variance (**Fig. 3C**). Separation according to genotype and treatment group was also evident, accounting for 3.6% (PC5) and 2.5% (PC6) of total variance (**Fig. 3C**). Expression of *bigmax*, the Mlx encoding gene, validated the sample genotypes and both Mondo and Mlx showed increased levels upon high fructose feeding (**Fig. 3D and Fig. S3C**). These results suggest that the previously reported feed-forward autoregulation of sugar responsive genes (Herman et al., 2012) is an evolutionary conserved mechanism.

Similar to the results in mouse, high fructose feeding induced robust gene expression changes (DEGs, adj.p < 0.05) in all studied regions (**Fig. 3E**). Mondo-Mlx appeared as the master regulator of fructose-responsive gene expression in the midgut, as only a minority of these fructose responsive genes showed a similar response in *mlx^1^* larvae (**Fig. 3E**). Mlx-dependent pathways that are regulated by fructose feeding, were mostly metabolism-related (**Fig. 3F**). Pathways of carbohydrate and amino acid metabolism were strongly overrepresented in the distal half of the midgut, consistent with regionalized midgut metabolism (**Fig. 3F**).

Analysis of individual Mondo-Mlx targets revealed both ubiquitous and regionalized expression. Transcription factor Cabut is a known direct target of Mondo-Mlx (Havula et al., 2013; Bartok et al., 2015) and its expression was induced by high fructose throughout the intestine in control animals but was very low and unresponsive to fructose in *mlx^1^* mutants (**Fig. 3G**). Its mammalian ortholog, *Klf10*, displays ubiquitous activation in all regions of the small intestine (**Fig. 3G**), consistent with evolutionary conservation of the regulatory axis between these sugar responsive TFs. On the other hand, expression of *Gs2*, encoding glutamine synthetase (Mattila et al., 2015), displayed highly regionalized expression in *Drosophila*, being strongly induced by fructose in a Mondo-Mlx-dependent manner in the distal midgut (LM4) (**Fig. 3H**). Also, its mammalian ortholog was fructose responsive throughout the small intestine, but like in flies, displayed a distally biased expression pattern (**Fig. 3H**). Consistent with the RNAseq data, whole mount staining of the *Drosophila* larval midguts show GS2 protein expressed in Mex1 positive enterocytes (Phillips and Thomas, 2006) of the LM4 region (**Fig. 3I**), and its expression in high fructose conditions was significantly reduced in *mlx^1^* animals (**Fig. 3J**). Collectively, our data is consistent with a conclusion that the intracellular sugar sensor Mondo/ChREBP-Mlx is a conserved mediator of the transcriptional response to high fructose in the intestines of *Drosophila* larva and mice. While Mondo-Mlx is active throughout the intestine (**Fig. 3A, E, G**), it also controls fructose-inducible expression of genes with a highly regionalized expression profile.

### Fructose feeding reduces the expression of cell cycle and DNA repair genes independently of Mondo-Mlx

A total of 2710 fructose-responsive DEGs were identified in control *Drosophila* larvae. Of these, 876 responded in similar way in control and *mlx^1^* mutants. Overrepresentation analysis revealed that most of these Mondo-Mlx-independent genes belong to pathways related to cell cycle and DNA repair (**Fig. 4A**). Genes belonging to these pathways include DNA polymerase delta complex, which is the major enzyme in eukaryotic chromosomal DNA replication, involved in many forms of DNA damage repair processes (Fuchs et al., 2021). DNA polymerase delta subunits 2 (*PolD2*) and its tethering sliding clamp PCNA were both downregulated in most of the intestinal regions (**Fig. 4B**). DNA repair process is assisted by Histone deacetylases (HDACs) and, in our samples, *HDAC1* expression was reduced in an Mlx-independent manner in all studied regions (**Fig. 4B**). Other genes that showed similar expression patterns were *Orc3*, the core subunit of Origin Recognition Complex (**Fig. 4B**); the initiation factor *Cdc6* (**Fig. 4B**) and *Mcm7,* the core subunit of the MCM2–7 helicase complex (**Fig. 4B**) (Faull et al., 2025).

**Figure 4.**
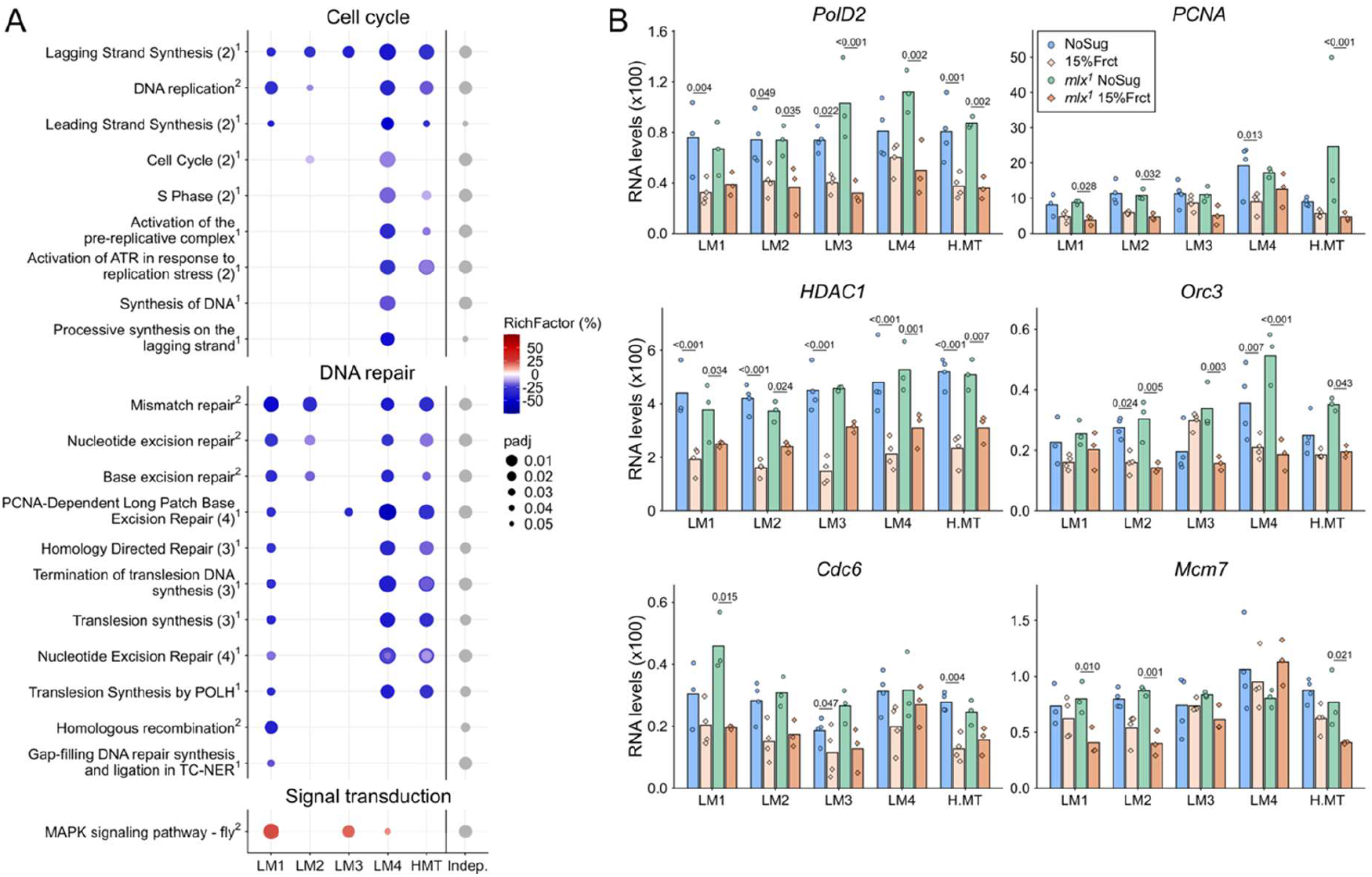
Mlx-independent response to fructose in the intestine consists of core cell cycle and DNA repair pathways. **A**) All regionally enriched Mlx-independent pathways resulted from over representation analysis (ORA), with *P* < 0.05 (dot size), adjusted for multiple comparison using the Benjamini–Hochberg correction. Pathway Richfactor resulting from upregulated DEGs are on a red scale and those resulting from downregulated DEGs were converted into negative values and shown on a blue scale. Databases used were: ^1^Reactome, ^2^KEGG. Enriched pathways with highly overlapping gene sets have been merged and total number of pathways in one row is reported within parenthesis, when merging has been applied. **B**) Expression levels of Mlx-independently fructose responsive example genes. P values were determined using DESeq2 and Benjamini-Hochberg correction. Adjusted p-values > 0.05 are not shown. N = 3-4 for all groups.

These results show that the Mondo-Mlx-independent genes in the midgut are mostly downregulated and related to specific cell cycle and DNA repair related processes.

### Evolutionary divergence and conservation of fructose-responsive intestinal gene expression

To systematically identify the common biological pathways responsive to high fructose feeding we performed a comparative analysis of the orthologs. Using Ensembl genome database we identified 11508 ortholog pairs in our data sets of which 1117 were differentially expressed upon fructose feeding and 636 were similarly regulated (**Fig. 5A**). Of these, 328 orthologs were upregulated, 308 were downregulated, and were distributed among all regions in both species (**Fig. 5A**). Over-representation analysis showed that the conserved upregulated genes were enriched in pathways related to glucose metabolism and its regulation, including insulin signaling and insulin resistance. In contrast, the downregulated genes formed two dominant functional clusters associated with RNA and protein metabolism (**Fig. 5B**), closely resembling the downregulated pathways identified in the mouse ileum (**Fig. 2E).**

**Figure 5.**
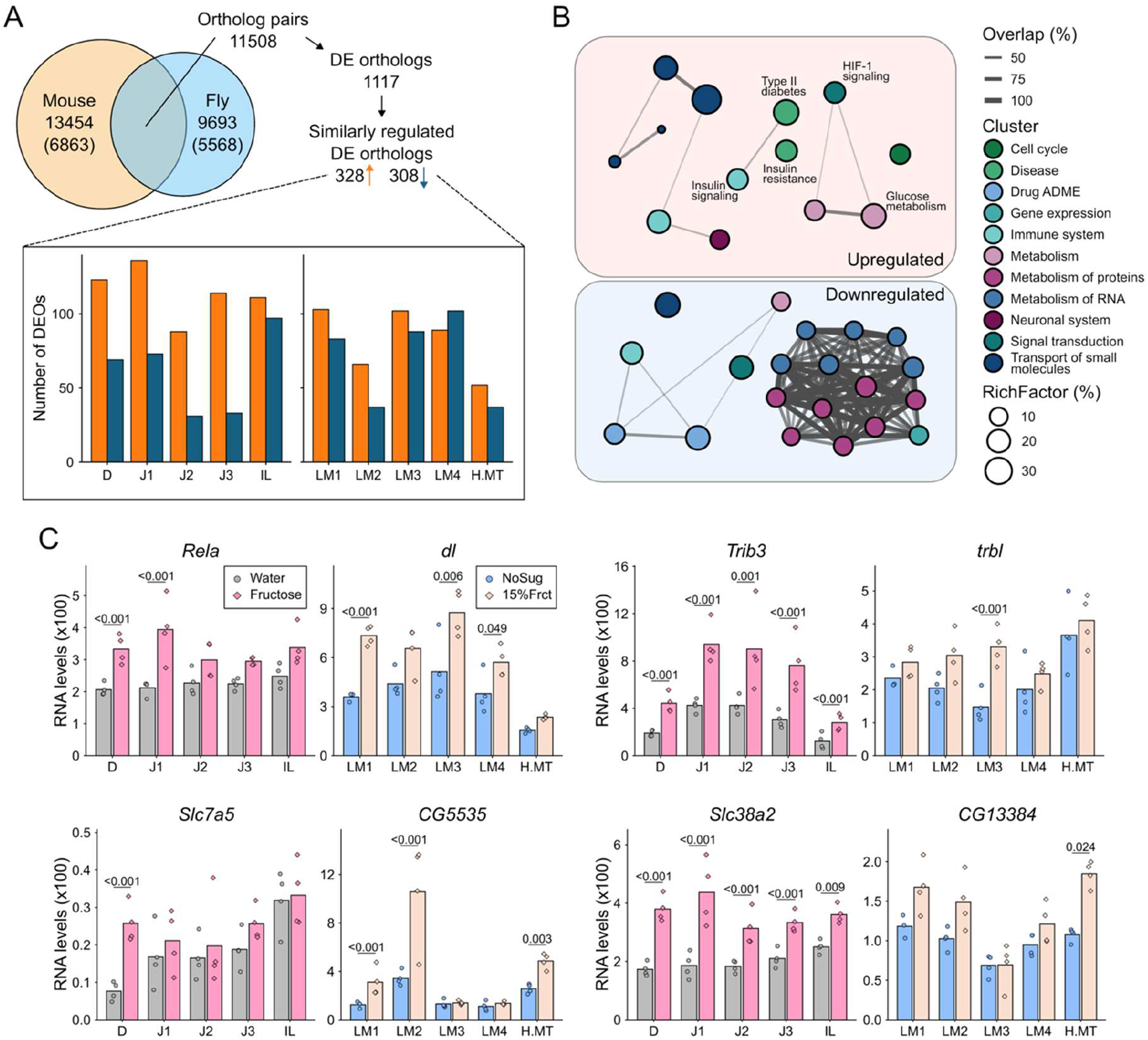
Fructose-responsive orthologs are enriched in conserved pathways related to glucose handling, nutrient transport, and cellular stress. **A)** Total number of protein coding genes and ortholog pairs mapped using the Ensembl database. Numbers of genes with at least one ortholog are shown in brackets. Regional distributions of genes with DE orthologs that are similarly regulated by fructose in both species are shown in the barplots. **B)** Enriched pathways with *P* < 0.05, adjusted for multiple comparison using the Benjamini–Hochberg procedure, resulting from analysis of DE orthologs. Each node represents an individual pathway, with node size representing Richfactor. Node colour represents the cluster to which the pathway is annotated to. Width of connecting lines represent the overlap of the sets of differentially expressed genes annotated to the pathways. **C)** Examples of genes with similar response to fructose in both species. P values were determined using DESeq2 and Benjamini-Hochberg correction. Adjusted p-values > 0.05 are not shown. N = 3-4 for all groups.

Several conserved fructose-responsive orthologs were associated with energy homeostasis and stress-responsive signaling. For example, *Rela*, encoding the p65 subunit of the NF-κB, a transcriptional mediator of diverse inflammatory and stress stimuli (Schottelius and Baldwin Jr., 1999), was upregulated regulated in response to fructose in a conserved manner (**Fig. 5C**). Consistent with being a direct target of NF-κB (Wu et al., 2003), also Trib3 was upregulated in both species (**Fig. 5C**). Together, these transcriptional changes indicate that acute fructose exposure engages conserved stress- and inflammation-related responses in the intestine of both species. Conserved responses were also observed among genes involved in the transport of small molecules. These included *Slc7a5*, encoding the large neutral amino acid transporter LAT1, and *Slc38a2*, encoding the sodium-coupled neutral amino acid transporter SNAT2 (**Fig. 5C**). Regulation of these transporters suggests that fructose exposure affects amino acid transport and cellular nutrient availability in both species.

Glucose metabolism emerged as a conserved fructose-responsive process when the mouse and Drosophila datasets were analyzed separately (**Fig. 1G** and **Fig. 3F**). Consistent with this enrichment, orthologs encoding key enzymes of glycolysis, including glucose-6-phosphate isomerase (Gpi), aldolase B (Aldob, also involved in fructolysis), pyruvate kinase (Pklr), as well as pyruvate dehydrogenase E1 component subunit beta (Pdhb), a component of the pyruvate dehydrogenase complex, were upregulated following high-fructose feeding in both species (**Fig. 6A, B**). In mammals, a substantial fraction of fructose can be converted to glucose and other metabolites in the intestine, thereby limiting systemic fructose exposure (**Fig. 6A**) (Öckerman and Lundborg, 1965; Jang et al., 2018). Consistent with this metabolic function, *Tkfc*, encoding triokinase/FMN cyclase, and *G6pc1*, encoding glucose-6-phosphatase catalytic subunit 1, were strongly upregulated throughout the mouse small intestine following fructose feeding. (**Fig. 6C**). In contrast, Tkfc and G6pc1 were not detectably expressed in the *Drosophila* midgut, suggesting that fructose is metabolized differently (**Fig. 6C**). In insects, glucose is primarily transported to the circulation as trehalose, whose synthesis involves trehalose-6-phosphate synthase (TPS) and trehalose-6-phosphate phosphatase (TPP), and occurs predominantly in the fat body (Matsuda et al., 2015). Consistent with the absence of a strong gluconeogenic response in the Drosophila midgut, expression of Tps and Tpp was low and largely unresponsive to fructose (**Fig. S4**).

**Figure 6.**
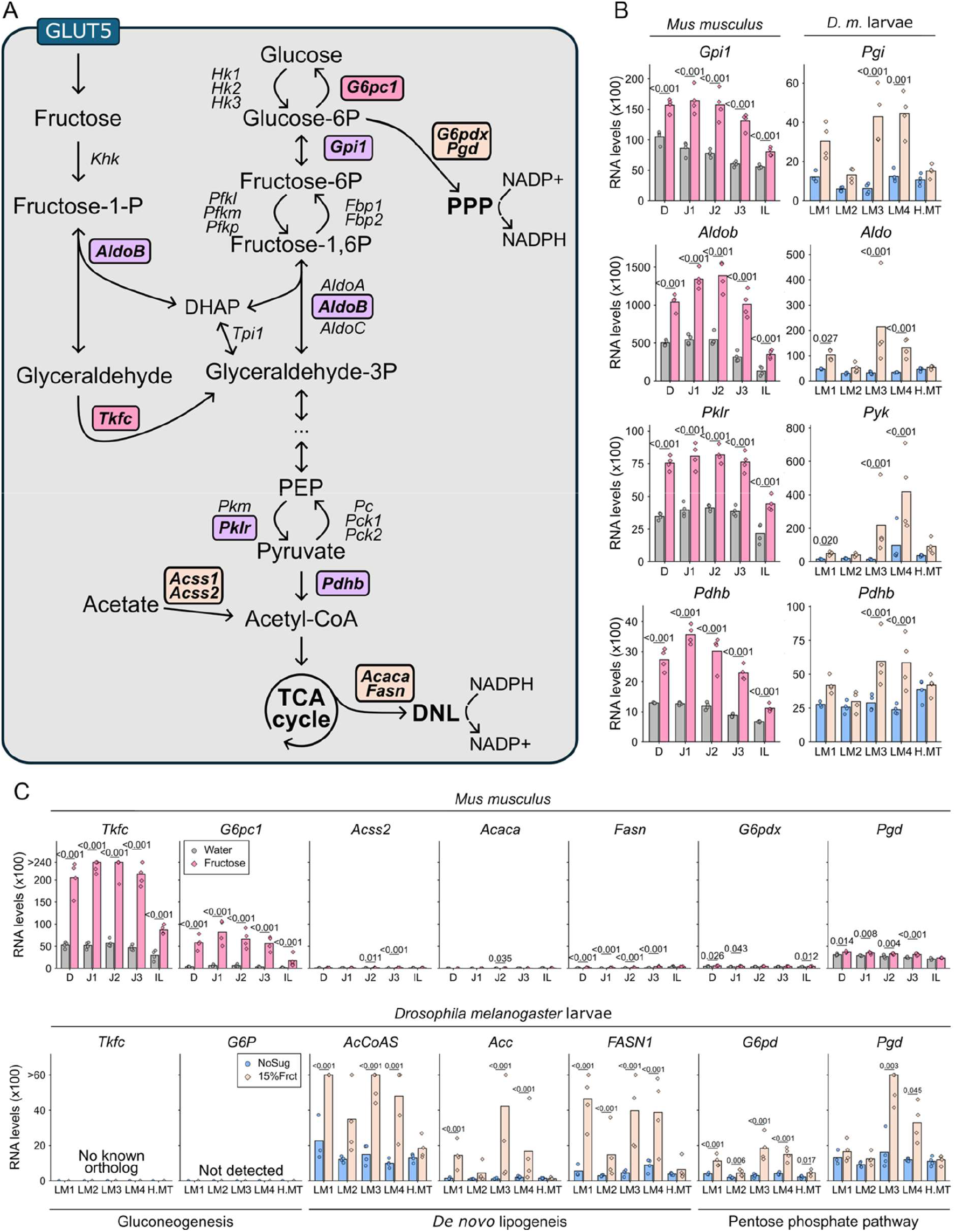
Transcriptional responses indicating species specific fructose processing in the intestine. **A**) Schematic of fructose metabolism highlighting its use in the gluconeogenesis or de novo lipogenesis (DNL) pathways, and the genes encoding the key enzymes. Intermediary metabolites labelled with abbreviations are: DHAP, Dihydroxyacetone phosphate; PEP, Phosphoenolpyruvate. **B**) Mouse and larvae RNA-seq data showing the expression changes for glycolytic genes and **C)** genes connected to fructose-specific gluconeogenic pathway (*Tkfc,* triokinase/FMN cyclase; *G6pc1*, Glucose-6-Phosphatase Catalytic Subunit), *de novo* lipogenesis (DNL) pathway (*Acss2*, Acetyl-coenzyme A synthetase; *Acaca*, acetyl-CoA carboxylase alpha; *Fasn*, fatty acid synthase) and pentose phosphate pathway (PPP; *G6pdx,* glucose-6-phosphate dehydrogenase*; Pgd,* phosphogluconate dehydrogenase) and their *Drosophila* orthologs. P values were determined using DESeq2 and Benjamini-Hochberg correction. Adjusted p-values > 0.05 are not shown. N = 3-4 for all groups.

In contrast, fructose strongly induced the expression of genes involved in *de novo* lipogenesis in the *Drosophila* midgut. *Acss2*, *Acaca* and *Fasn*, encoding Acetyl-coenzyme A synthetase, acetyl-CoA carboxylase and fatty acid synthase, respectively, were strongly upregulated following fructose feeding, whereas their mouse orthologs showed low expression and little or no fructose response in the small intestine (**Fig. 6C**). Consistent with increased lipogenic activity, the pentose phosphate pathway, which provides NADPH required for fatty acid synthesis, were enriched in Drosophila but not in mouse (**Fig. 6A, C**). Collectively, both species activate metabolic gene expression in the intestine to enable conversion of fructose into less harmful metabolites (fructose shielding), but they likely utilize distinct metabolic strategies to achieve this.

Collectively, the comparative analysis revealed both conserved and divergent features of the intestinal response to acute fructose exposure. Conserved responses included activation of glucose metabolism, nutrient transport, and stress- and signaling-related pathways, whereas the regulation of downstream metabolic pathways differed substantially between species. These findings suggest that the intestinal capacity to metabolically process fructose is evolutionarily conserved, while the metabolic pathways used to accommodate fructose exposure have diverged in accordance with species-specific carbohydrate physiology.

## DISCUSSION

The emerging understanding of the metabolic organization of the small intestine has opened a new perspective on how to study nutritional challenges and the molecular changes that underlie them (Zwick et al., 2026, 2024). Fructose has been shown to act strongly on selected genes in the small intestine, but its genome-wide and region-specific effects have thus far remained uncharacterized (Kim et al., 2017). The data generated in this study offers several new insights into gene regulatory responses to fructose, including its spatial organization and evolution. We observed that acute fructose feeding triggers a robust, intestine-wide transcriptional response in both mice and *Drosophila*, including expected changes in metabolic genes, but also signs of stress and immune pathway activation. The intracellular sugar sensor Mondo-Mlx emerged as a central regulator of fructose-responsive intestinal gene regulation in Drosophila, and its conserved targets were strongly fructose responsive in mouse villi. Regional analysis revealed that while many of the Mondo/ChREBP-Mlx targets were controlled mostly ubiquitously, several metabolic and stress-related pathways displayed spatial expression patterns. Cross-species comparison revealed that although a core set of orthologous genes involved in glucose metabolism, nutrient transport, and stress responses show conserved regulation, mice and flies rely on distinct metabolic strategies for fructose shielding (glucose-6-phosphatase-based versus fatty-acid-synthesis-based pathways). Together, these findings imply an ancestral role for intestinal gene regulation in controlling fructose metabolism in order to shield other organs, but the specific metabolic strategies involved appear to have diverged during evolution.

ChREBP/Mondo-Mlx emerged as a conserved central regulator of the intestinal response to high fructose. In mice, canonical ChREBP-Mlx target genes, including *Mlxipl, Txnip*, and *Slc2a5*, were induced throughout the small intestine, although to a lesser extent in the ileum, indicating that ChREBP-Mlx activation is not restricted to a discrete intestinal domain. This organ-wide activity is consistent with the finding that intestinal ChREBP is required for mice to tolerate a high-fructose diet (Kim et al., 2017). In Drosophila, Mondo-Mlx controlled the fructose response of several hundred genes across all intestinal regions, further supporting its role in regulating a broad transcriptional program that facilitates tolerance to high dietary fructose. ChREBP/Mondo-Mlx is an established regulator of intracellular sugar sensing that acts together with other transcription factors (Dentin et al., 2004; Havula et al., 2013; Kibbe et al., 2013; Bartok et al., 2015; Mattila et al., 2015; Kokki et al., 2021; van den Berg et al., 2023; Tsukamoto et al., 2024; Kokki et al., 2026). Cooperation with regionally restricted factors may help explain the spatially distinct responses observed here. For example, cholesterol biosynthesis was induced in both the duodenum and ileum, suggesting potential cooperation between Srebp2 and ChREBP-Mlx, as described in the liver (Zhang et al., 2017). In contrast, HSF1-associated heat-shock responses were biased toward the proximal intestine, whereas the ileum showed stronger JAK/STAT- and interferon-associated inflammatory programs. Thus, ChREBP-Mlx-dependent metabolic sensing appears to be integrated with regionally distinct stress- and immune-responsive transcriptional networks.

The marked differences between proximal and distal small intestine provide insight into how the tissue accommodates to high fructose ingestion. The intestine acts as a protective barrier against systemic fructose exposure by converting a substantial fraction of ingested fructose into glucose and organic acids before portal entry (Bismut et al., 1993; Jang et al., 2018, 2020; Zhao et al., 2020; Herman and Birnbaum, 2021). The proximal intestine appears to have a greater capacity for fructose processing, but this may come at the cost of increased cellular stress. Fructose uptake leads to rapid, poorly regulated conversion to fructose-1-phosphate, which can deplete cellular ATP and induce stress (Mäenpää et al., 1968; Herman and Birnbaum, 2021). Consistent with this, we observed induction of heat-shock-responsive chaperones in the proximal intestine, including Hsph1, which can suppress aggregation of denatured proteins under severe cellular stress associated with ATP depletion (Yamagishi et al., 2003).

In contrast, the lower expression of key fructose-shielding genes such as *Tkfc* and *G6pc1* indicates that the ileum has a more limited capacity to process fructose may be particularly vulnerable to fructose-induced stress. Consistent with this possibility, JAK/STAT-, interferon-, IRF7-, and Jun kinase-associated pathways were activated in the ileum. The pro-inflammatory effects of high-fructose diets have been demonstrated in experimental models of colitis and have been linked to changes in the gut microbiota and disruption of the intestinal epithelial barrier (Kawabata et al., 2019; Montrose et al., 2021). Studies in humans, primates, and mice further indicate that fructose-induced disruption of the intestinal barrier can contribute to endotoxemia and liver fat accumulation (Bergheim et al., 2008; Jin et al., 2014; Kavanagh et al., 2013; Todoric et al., 2020). Our findings suggest that the ileum may represent an important site of this response, where limited fructose-shielding capacity could promote cellular stress and activation of inflammatory pathways. Thus, intestinal fructose metabolism may serve not only to protect peripheral organs from fructose exposure but also to protect the intestinal tissue itself from metabolic and immune consequences of fructose overload.

Comparison of regional transcriptomes from mice and Drosophila revealed both conserved and species-specific features of intestinal fructose responses. Mondo-Mlx, like mammalian ChREBP-Mlx, functions as a broadly active intestinal sugar sensor while regulating targets with either ubiquitous or strongly regionalized expression. Genome-wide ortholog analysis identified conserved responses, including genes involved in glucose metabolism, nutrient transport, and stress responses. Individual examples, such as *Drosophila cabut* and *GS2* and their mammalian orthologues *Klf10* and *Glul*, further illustrate this conservation. Cabut/Klf10, for example, integrates sugar metabolism and circadian regulation in both species, supporting functional conservation of this regulatory connection (Bartok et al., 2015; Ruberto et al., 2021). At the same time, the downstream metabolic strategies for fructose shielding have diverged. In mice, fructose-responsive intestinal gene expression was associated with glucose production, including induction of pathways involving glucose-6-phosphatase. In Drosophila, by contrast, Mondo-Mlx strongly induced genes involved in de novo lipogenesis, consistent with the prominent role of intestinal lipogenesis in this species (Palm et al., 2012). These findings suggest that the ancestral function of the Mondo/ChREBP-Mlx system as a sensor of intracellular sugar availability has been conserved, whereas its downstream metabolic outputs have adapted to the distinct physiological organization and nutrient-handling strategies of mammals and insects.

## MATERIALS AND METHODS

### Mouse Experiments

Animal procedures were approved by the Finnish Supervisory Agency, license number ESAVI/11519/2023. We used 12–13-week-old C57BL/6J mice due to their established sensitivity to metabolic diseases, particularly in dietary studies (Kleinert et al., 2018). Only male mice were included due to their stronger response to dietary challenges, the focused scope of the study, and available resources. Mice were fasted for 5 hours prior to fructose or water administration by oral gavage and killed two hours later, by gradual C0_2_ intoxication followed by cervical dislocation. Fructose concentration (4g/kg, diluted in drinking water at 30% concentration) was selected according to the study by Kim et al. 2017 (Kim et al., 2017), but similar concentrations were used also in other studies (Goncalves et al., 2019; Taylor et al., 2021). Small intestine was removed and flushed with cold PBS, and cut into five segments as recently reported (Zwick et al., 2024). Each segment was cut open and further cleaned with a brush soaked in cold PBS. Villi were collected by scraping the luminal side of the intestine with a clean microscopy glass, without excessive pressure to minimize the intestinal crypts disruption collection, and snap frozen in liquid nitrogen (**Figure 1A**). Only one biological replicate in each group was prepared per day to minimize technical errors and improve the reproducibility of the results.

### Drosophila melanogaster larvae experiments

Mlx null mutant *Drosophila melanogaster* flies and their matched controls were previously described (Havula et al., 2013). Stocks were kept under controlled conditions (18°C with a consistent 12 h light-dark circadian cycle) or between 21-25°C, fed a basic laboratory diet composed of 6.5% malt (w/v), 3.2% semolina (w/v), 1.8% dry baker’s yeast (w/v), 0.6% agar (w/v), 0.7% propionic acid (v/v), and 2.5% nipagine (methyl paraben) (v/v). For larvae collection, flies were placed in custom egg laying chambers equipped with 5-cm petri dishes containing 33.3% apple juice, 1.75% agar, 2.5% sugar, 2.0% nipagin, and sprinkled with few crumbs of yeast. First instar larvae were picked and transferred on a chemically defined diet (Piper et al., 2017), and allowed to grow until they reached early third instar stage before they were transferred on the same diet without sugar (NoSug) or containing 15% fructose (Fru15%) for six hours. For protein expression imaging, larvae were fed the experimental diets for 24 hours. Gut fragments were cut with small scissors under a stereomicroscope as depicted in **Fig. 3B**. Alkaline region was visualized by addition of 0.2% mCresol in the experimental diets (Overend et al., 2016). Each biological replicate contained gut fragments from 5 different larvae collected in RNA lysis buffer and snap frozen in liquid nitrogen. Only one biological replicate in each group was prepared per day to minimize technical errors and improve the reproducibility of the results.

### RNA-sequencing

Mouse samples RNA extraction was done using NucleoSpin RNA extraction kit (Macherey-Nagel), RNA integrity was tested using Bioanalyzer (Agilent), and the library was made with Illumina Stranded TruSeq mRNA Library Prep Kit. Single strand 75 bp paired-end RNA-sequencing was performed using AVITI High Output (Element Biosciences).

*Drosophila* samples RNA extraction was done using NucleoSpin RNA XS extraction kit (Macherey-Nagel), RNA integrity was tested using Bioanalyzer (Agilent RNA 6000 Nano), and the library was made with Illumina TruSeq Stranded mRNA LT Library Prep Kit. Single strand 75 bp single-end RNA-sequencing was performed using Illumina NextSeq 500.

### Immunofluorescence imaging

For immunofluorescence staining, feeding larvae were partially dissected in saline (0.9 % NaCl) to expose visceral tissues and transferred to cold 4% formaldehyde in PBS (Thermo Fisher #28908) for overnight fixation in cold with gentle rocking. Fixed larval tissue was permeabilized, blocked and stained in PBS, 1 % BSA, 0.1 % TritonX-100 and washings were done in PBS, 0.1 % TritonX-100. Primary antibodies (1:500), phalloidin (1:500, Invitrogen, cat# A12380) and secondary antibodies (1:1000, Invitrogen/Molecular Probes Cat# A-11008, RRID: AB_143165) were diluted in staining buffer and incubated with the samples overnight in cold with gentle rocking. Samples were incubated in Vectashield with DAPI (Vectashield #H-1200) overnight in cold, followed by full dissection and flat mounting with a custom 60 µm spacer (Scotch Magic Tape 810). Anti-Mlx antibodies were generated in house and previously reported (Havula et al., 2013). Briefly, full-length *Drosophila mlx* cDNA was cloned into pGEX-4T2 and recombinant GST-Mlx was purified using Glutathione-agarose (Sigma). Anti-Mlx antiserum was raised by immunizing a guinea pig (Storkbio Ltd). Purified Mouse Anti-Glutamine Synthetase antibodies (RRID: AB_397879) were purchased from BD Transduction Laboratories. Enterocytes were visualized using a transgenic line expressing Mex1-GFP positive enterocytes (Phillips and Thomas, 2006). Images were acquired with Aurox Clarity LFC HS and Leica SP8 laser confocal microscope and processed using Fiji/ImageJ.

### Analysis of bulk RNA-sequencing data

The bulk RNA-seq datasets were processed using nf-core/rnaseq pipeline (v.3.22.2) from the nf-core collection of workflows, utilizing environments from the Bioconda and Biocontainers projects. The pipeline was executed with Nextflow (v. 25.10.2). FastQC (v.0.12.1) was used for quality metrics and fastp (v. 0.24) was used for read trimming with standard settings. The reads were mapped using STAR (v. 2.7.11b) to reference genome dmel_r6.66 and GRCm39 for *Drosophila* and mouse respectively. The mapping was quantified with Salmon (v. 1.10.3). NCBI RefSeq (v. 236) was used for retrieving the missing gene biotype annotations for *Drosophila* dataset. The raw sequencing data is available on GEO, accession number GSE349771 and GSE349772 for *Drosophila* and mouse respectively.

### Preprocessing of single-cell RNA-sequencing data

Single-cell RNA-sequencing data was obtained from previously published study (Zwick et al., 2024). The data had already been assessed for quality control, SCTransform normalization and Seurat (v.5.5.0) integration as reported in the source study. The high-fat samples and cell types other than enterocytes were excluded from all downstream analyses and the 15 segments were merged to correspond to the five regions used in this study by combining segments 1-2, 3-5, 6-10, 11-12 and 13-15 to form D, J1, J2, J3 and IL respectively based on the Pearson correlation of the segments’ expression profiles and known relative lengths of the regions. The data was converted into pseudobulk by summing the raw counts of each unique combination of region, diet and mouse, to produce total of 30 pseudobulk samples.

### Differential expression analysis

Differential expression analysis was conducted with DESeq2 (v. 1.46.0) in R (v. 4.4.2)/Bioconductor (v.3.20). Genes with minimum of ten reads in minimum of four samples in our bulk datasets and minimum of three samples in single-cell derived pseudobulk dataset were retained. Size factors were estimated using “poscounts” and dispersions were fit using “parametric” methods. For plotting gene expression levels, the bulk RNAseq counts were further normalized by dividing the counts by Salmon derived per-sample transcript lengths. For plotting single-cell derived pseudobulk counts were converted into counts per million (CPM). No transcript length normalization was applied due to the nature of Unique Molecular Identifier (UMI)-based capture. Principal component analysis (PCA) was conducted for all filter passing genes following variance-stabilizing transformation. The design formulas contained all variables and their interaction terms. Non-protein coding genes were included in differential expression analysis but filtered out from the data used for downstream analysis. Differential expression analysis was conducted using Wald test and p-values were adjusted using Benjamini-Hochberg method. Fructose-induced changes were measured using the diet coefficient (mouse and control genotype in the Drosophila dataset) or the diet + genotype:diet coefficients (*mlx* null). The Mlx-dependence of the fructose response was tested using the genotype:diet interaction coefficient. Genes with adjusted p-value less or equal to 0.05 and absolute log2 fold change greater or equal to 0.5 were considered differentially expressed. For *Drosophila* data the genes were classified as mlx dependent if the dependency and the fructose-induced changes in control genotype reached the above-mentioned cutoffs for differential expression and no fructose-induced changes or opposite change was detected in *mlx-*null genotype. The DESeq2 Wald test derived adjusted p-values were used in the expression level plots for all datasets.

### Assessment of cell type composition

Relative proportions of the intestinal cell types in the bulk RNAseq samples were estimated with Multi-Subject Single Cell deconvolution (MuSiC, v. 1.0.0) (Wang et al., 2019). Previously published single cell RNAseq dataset from mouse small intestine (Chen et al., 2025) was used as reference dataset. Quality thresholds were matched with the original study and replicates Control_1, Fruc_2 and HFHS_1 were removed from the reference due to missing cell types and low overall cell counts. Cell types were called from the author-provided metadata. The raw Salmon-derived readcounts were used as input from the bulk RNAseq. MuSiC was run in R/Bioconductor with default parameters.

### Ortholog mapping

Orthologs were retrieved bidirectionally from Ensembl database (v. 115). In the case of multiple orthologs per gene, all pairs were kept. Pairs with orthologs expressed in both species were kept for downstream analysis. Ortholog pairs were considered similarly expressed if the high-fructose diet induced significant differential gene expression in the same direction in at least one region in both mouse and *Drosophila* control data.

### Pathway enrichment analysis and enrichment maps

Over-representation analysis was performed with clusterProfiler (v. 4.14.6) in R (v. 4.4.2)/Bioconductor (v.3.20) using Kyoto Encyclopedia of Genes and Genomes (KEGG, v. 119.0) and Reactome (v. 97) databases. All protein coding genes that reach the expression level cutoff in our datasets were used as background for the analysis within species. For differentially expressed ortholog pairs, the mouse counterparts were used as input and all protein coding genes with an ortholog that was expressed also in *Drosophila* were used as background. Pathways with adjusted p-value less or equal to 0.05 with Benjamini-Hochberg correction were considered enriched.

From *Drosophila* dataset mlx-dependent pathways were identified by performing over-representation analysis for mlx-dependent genes with DEGs from control group as background, both pooled across regions by keeping genes that were dependent/DEG in any of the regions. The resulting pathways were filtered to the overlap with the fructose responsive pathways within control genotype in any region prior to p-value adjustment by Benjamini-Hochberg correction. For dependency a cutoff of 0.10 was used for the adjusted p-value to consider the pathway mlx-dependent.

The Mlx-independence of pathways was assessed regionally by performing permutation test for all pathways enriched with adjusted p-value of 0.25 or less in both genotype’s fructose response over-representation analysis. For each pathway, the mean of the DESeq2 log2 fold change (LFC) for the diet-genotype interaction contrast, computed over the union of the pathway’s leading-edge genes was used as test statistic, while the interaction LFCs of all genes differentially expressed upon fructose feeding in either of the genotypes were used as null distribution. 10 000 permutations of the size of the union of the leading edges were drawn without replacement and the interaction LFC means were calculated. Lower-tail p-value was calculated by dividing one plus the number of permutations where the interaction LFC mean was smaller than the observed by one plus the number of permutations. P-values were adjusted for multiple testing using Benjamini-Hochberg correction. If the same pathway was enriched within both genotypes in multiple regions, the most significant adjusted p-value was reported for the Mlx-independence. Pathways found independent with adjusted p-value less or equal to 0.05 were plotted together with control genotype’s enrichment results.

For plotting, pathways with Jaccard similarity of 0.8 or greater of their contributing sets of differentially expressed genes were merged in order to avoid overemphasizing redundant results. The names of the merged pathways were manually assessed taking into an account the degree of RichFactors of participating pathways. The number of pathways merged into one has been stated at the end of the pathway name within parenthesis. Furthermore, the ancestral class for each pathway from Reactome database and the category for each pathway from KEGG was used as cluster, with the exception of merging KEGG categories “Human Diseases”, “Environmental Information Processing”, “Genetic Information Processing”, “Organismal Systems” and “Cellular Processes” into Reactome classes “Disease”, “Signal transduction”, “Gene expression”, “Immune system” and “Cell cycle”, respectively. Missing clusters and those for merged pathways resulting from multiple clusters were manually assessed. Richfactors of pathways resulting from downregulated DEGs were converted into negative values. For enrichment maps the overlaps were calculated using Jaccard similarity of the differentially expressed genes of the pathway. Maps were plotted using igraph (v. 2.2.2) and ggraph (v. 2.2.2) packages in R (v. 4.4.2)/Bioconductor (v.3.20).

### Data visualization

Graphs were generated in R (v. 4.4.2) using packages ggplot2 (v.4.0.2), ComplexUpset (v.1.3.3), igraph (v.2.2.2) and ggraph (v.2.2.2). Inkscape was used for creating the figure layout and editing the plots.

## ACKNOWLEDGMENTS

We thank Heini Lassila for technical assistance, and Bohdana Rovenko, Onur Deniz, Jaakko Mattila and Rita Cerejeira Matos for their advice regarding the *Drosophila* experiments, and several other members of Hietakangas laboratory for their support and inspiring scientific discussions. We want to also thank Ophir Klein and Dario Boffelli for their assistance in providing and clarifying the files of the high-carbohydrate dataset. The study was facilitated by the University of Helsinki DNA Sequencing and Genomics Laboratory, Helsinki *Drosophila* core facility (Hi-Fly), Laboratory Animal Centre (LAC) and the Light microscopy unit (LMU), with support from Biocenter Finland and Helsinki Institute of Life Science; and by CSC – IT Center for Science, Finland, for providing computational resources.

This work was supported by the Research Council of Finland (grant # 336197, 372024 and 374086 to V. H.), Sigrid Juselius Foundation to V. H., Novo Nordisk Foundation (grant # NNF22OC007841 and NNF25OC0105265 to V. H.)

---

**Supplementary Figure 1.**
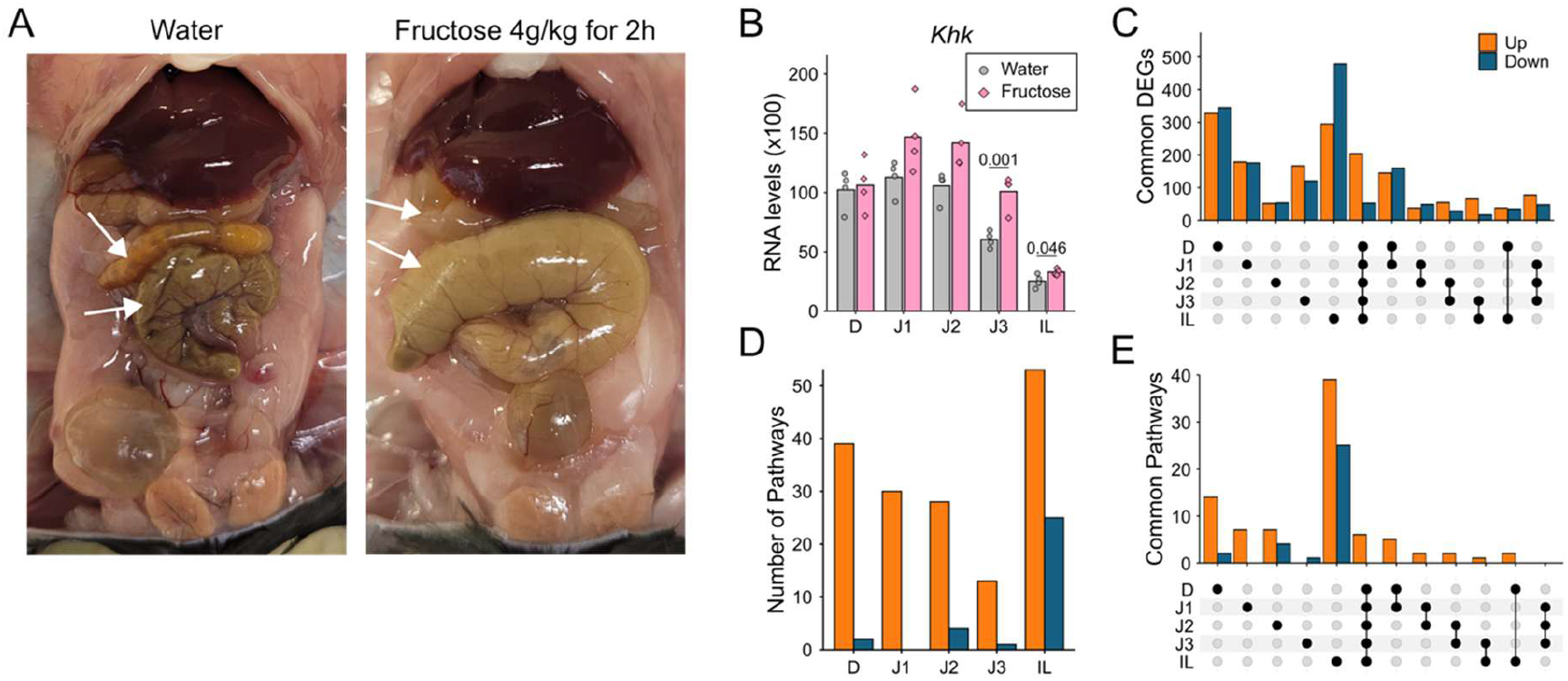
High fructose feeding leads to intestinal bloating and a strong transcriptional response in mouse small intestinal villi. **A**) Representative images showing bloating of fructose-fed mice observed during sample collection. Mice were fasted for five hours before the oral gavage. Small intestine and cecum are indicated with arrows. **B**) Expression levels of ketohexokinase (KHK). **C**) Total number of differentially expressed genes (DEGs) from Fig. 1F distributed as region-specific or region shared. **D**) Total number of enriched pathways in each region resulted from over representation analysis (ORA). Upregulated pathways resulted from upregulated DEGs and downregulated pathways from downregulated DEGs. **E**) Total number of enriched pathways from panel **D** distributed as region-specific or region shared.

**Figure S2.**
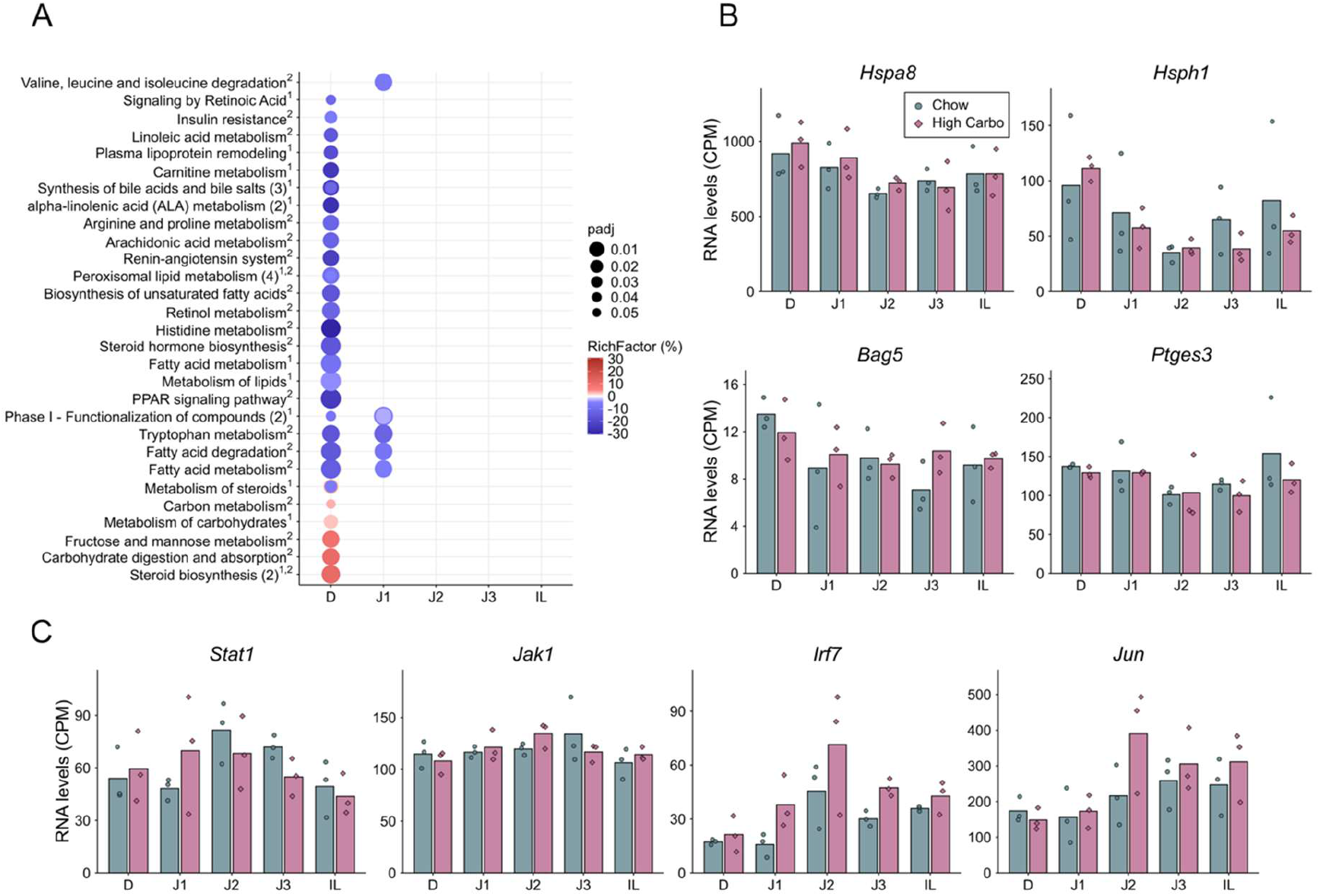
Acute high fructose, but not chronic high carbohydrate feeding, triggers stress- and immune-related gene responses in a region-specific manner. **A)** All regionally enriched pathways upon chronic high carbohydrate feeding resulted from over representation analysis (ORA), with *P* < 0.05 (dot size), adjusted for multiple comparison using the Benjamini–Hochberg correction. Pathway Richfactor resulting from upregulated DEGs are on a red scale and those resulting from downregulated DEGs were converted into negative values and shown on a blue scale. Databases used were: ^1^Reactome, ^2^KEGG. Enriched pathways with highly overlapping gene sets have been merged and total number of pathways in one row is reported within parenthesis, when merging has been applied. B) Expression levels of example genes related to heat shock response from Fig 2D. C) Expression levels of example genes related to stress and inflammation from Fig 2F. Statistical significance for B and C, was tested with DESeq2 and Benjamini-Hochberg correction. Adjusted p-values > 0.05 are not shown. N = 3 for all groups.

**Figure S3.**
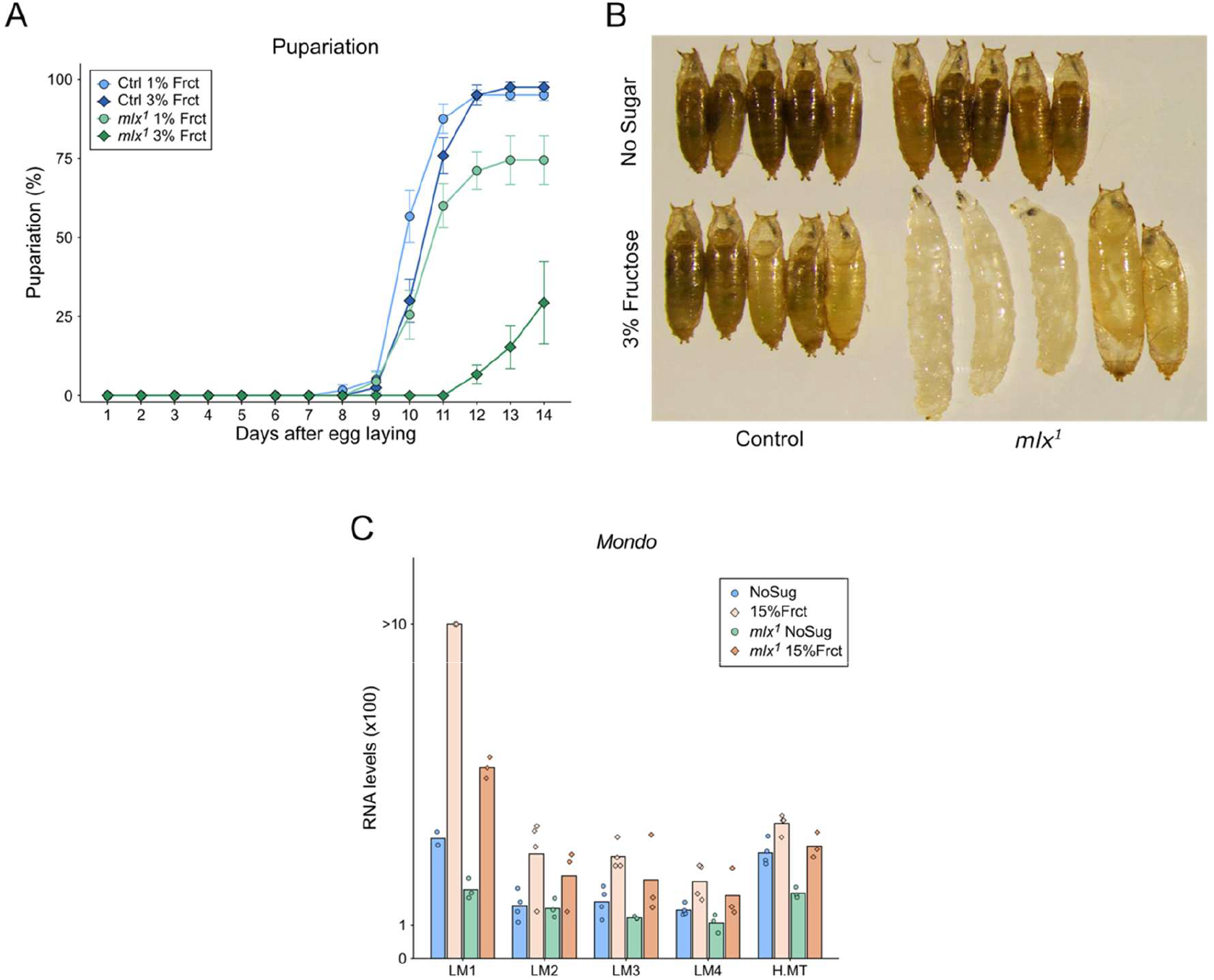
*Mlx*-null larvae are highly sensitive to dietary fructose. **A**) Pupariation measurements in Control and *mlx*-null larvae (*mlx^1^*) fed a holidic diet containing 1% or 3% fructose. **B**) Pupae images showing delayed and aberrant pupae formation in *mlx*-null larvae fed 3% fructose. **C**) Expression levels of Mlx dimerisation partner Mondo. In control genotype LM1 region, one outlier sample from each diet was removed from the plot due to over 8-fold higher expression compared to their respective replicates. Statistical significance was tested with DESeq2 and Benjamini-Hochberg correction. Adjusted p-values > 0.05 are not shown. N = 3-4 for all groups.

**Figure S4.**
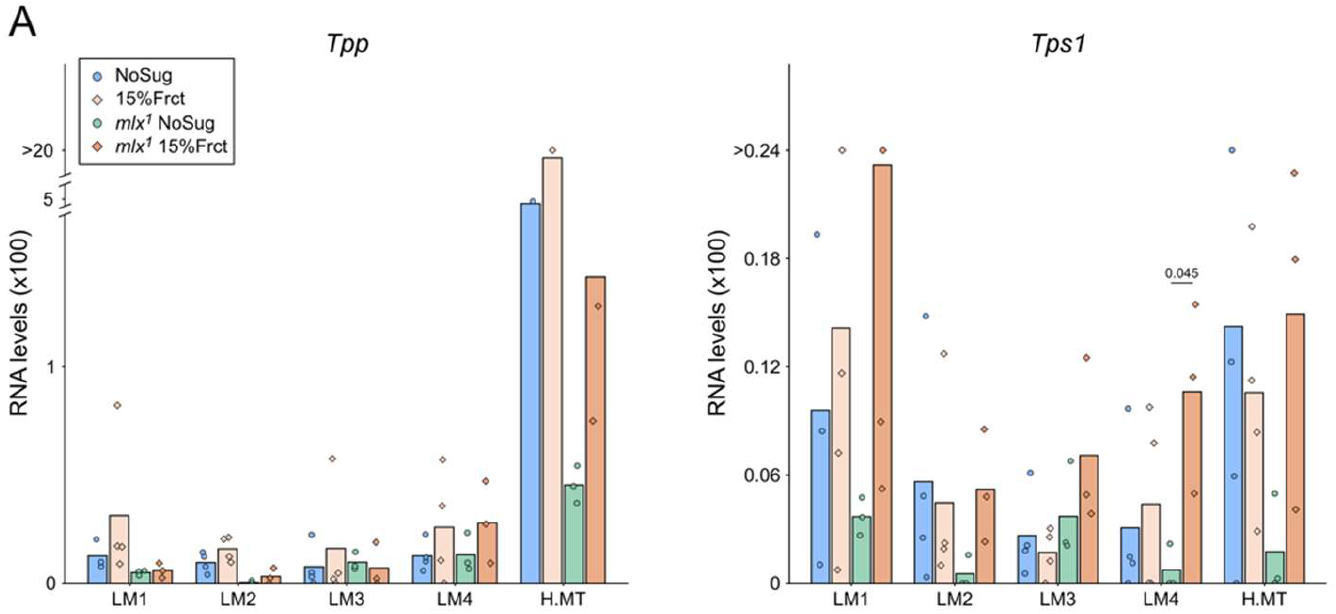
Trehalose biosynthetic genes are not activated by fructose in the midgut. **A**) Expression levels of genes coding key enzymes of trehalose producing pathway. P values were determined using DESeq2 and Benjamini-Hochberg correction. Adjusted p-values > 0.05 are not shown. N = 3-4 for all groups.

## REFERENCES

Adolph, T.E., Tilg, H., 2024. Western diets and chronic diseases. Nat Med 30, 2133–2147. 10.1038/s41591-024-03165-6

Bartok, O., Teesalu, M., Ashwall-Fluss, R., Pandey, V., Hanan, M., Rovenko, B.M., Poukkula, M., Havula, E., Moussaieff, A., Vodala, S., Nahmias, Y., Kadener, S., Hietakangas, V., 2015. The transcription factor Cabut coordinates energy metabolism and the circadian clock in response to sugar sensing. EMBO J 34, 1538–1553. 10.15252/embj.201591385

Bates, M.D., Erwin, C.R., Sanford, L.P., Wiginton, D., Bezerra, J.A., Schatzman, L.C., Jegga, A.G., Ley-Ebert, C., Williams, S.S., Steinbrecher, K.A., Warner, B.W., Cohen, M.B., Aronow, B.J., 2002. Novel genes and functional relationships in the adult mouse gastrointestinal tract identified by microarray analysis. Gastroenterology 122, 1467–1482. 10.1053/gast.2002.32975

Bismut, H., Hers, H.-G., Van Schaftingen, E., 1993. Conversion of fructose to glucose in the rabbit small intestine. European Journal of Biochemistry 213, 721–726. 10.1111/j.1432-1033.1993.tb17812.x

Cadena Del Castillo, C.E., Deniz, O., van Geest, F., Rosseels, L., Stockmans, I., Robciuc, M., Carpentier, S., Wölnerhanssen, B.K., Meyer-Gerspach, A.C., Peterli, R., Hietakangas, V., Shimobayashi, M., 2025. MLX phosphorylation stabilizes the ChREBP-MLX heterotetramer on tandem E-boxes to control carbohydrate and lipid metabolism. Sci Adv 11, eadt4548. 10.1126/sciadv.adt4548

Chen, Y.-W., Ahn, I.S., Wang, S.S.-M., Majid, S., Diamante, G., Cely, I., Zhang, G., Cabanayan, A., Komzyuk, S., Bonnett, J., Arneson, D., Yang, X., 2025. Multitissue single-cell analysis reveals differential cellular and molecular sensitivity between fructose and high-fat high-sucrose diets. Cell Reports 44. 10.1016/j.celrep.2025.115690

Cláudio, N., Dalet, A., Gatti, E., Pierre, P., 2013. Mapping the crossroads of immune activation and cellular stress response pathways. EMBO J 32, 1214–1224. 10.1038/emboj.2013.80

Costa-Mattioli, M., Walter, P., 2020. The integrated stress response: From mechanism to disease. Science 368, eaat5314. 10.1126/science.aat5314

Faull, S.V., Barbon, M., Mossler, A., Yuan, Z., Bai, L., Reuter, L.M., Riera, A., Winkler, C., Magdalou, I., Peach, M., Li, H., Speck, C., 2025. MCM2-7 ring closure involves the Mcm5 C-terminus and triggers Mcm4 ATP hydrolysis. Nat Commun 16, 14. 10.1038/s41467-024-55479-1

Febbraio, M.A., Karin, M., 2021. “Sweet death”: Fructose as a metabolic toxin that targets the gut-liver axis. Cell Metabolism 33, 2316–2328. 10.1016/j.cmet.2021.09.004

Fuchs, J., Cheblal, A., Gasser, S.M., 2021. Underappreciated Roles of DNA Polymerase δ in Replication Stress Survival. Trends in Genetics 37, 476–487. 10.1016/j.tig.2020.12.003

Goncalves, M.D., Lu, C., Tutnauer, J., Hartman, T.E., Hwang, S.-K., Murphy, C.J., Pauli, C., Morris, R., Taylor, S., Bosch, K., Yang, S., Wang, Y., Van Riper, J., Lekaye, H.C., Roper, J., Kim, Y., Chen, Q., Gross, S.S., Rhee, K.Y., Cantley, L.C., Yun, J., 2019. High-fructose corn syrup enhances intestinal tumor growth in mice. Science 363, 1345–1349. 10.1126/science.aat8515

Havula, E., Hietakangas, V., 2018. Sugar sensing by ChREBP/Mondo-Mlx-new insight into downstream regulatory networks and integration of nutrient-derived signals. Curr Opin Cell Biol 51, 89–96. 10.1016/j.ceb.2017.12.007

Havula, E., Hietakangas, V., 2012. Glucose sensing by ChREBP/MondoA–Mlx transcription factors. Seminars in Cell & Developmental Biology, Nutrient Sensing Pancreas Development 23, 640–647. 10.1016/j.semcdb.2012.02.007

Havula, E., Teesalu, M., Hyötyläinen, T., Seppälä, H., Hasygar, K., Auvinen, P., Orešič, M., Sandmann, T., Hietakangas, V., 2013. Mondo/ChREBP-Mlx-regulated transcriptional network is essential for dietary sugar tolerance in Drosophila. PLoS Genet 9, e1003438. 10.1371/journal.pgen.1003438

Herman, M.A., Birnbaum, M.J., 2021. Molecular aspects of fructose metabolism and metabolic disease. Cell Metabolism 33, 2329–2354. 10.1016/j.cmet.2021.09.010

Herman, M.A., Peroni, O.D., Villoria, J., Schön, M.R., Abumrad, N.A., Blüher, M., Klein, S., Kahn, B.B., 2012. A novel ChREBP isoform in adipose tissue regulates systemic glucose metabolism. Nature 484, 333–338. 10.1038/nature10986

Iizuka, K., Bruick, R.K., Liang, G., Horton, J.D., Uyeda, K., 2004. Deficiency of carbohydrate response element-binding protein (ChREBP) reduces lipogenesis as well as glycolysis. Proceedings of the National Academy of Sciences 101, 7281–7286. 10.1073/pnas.0401516101

Jang, C., Hui, S., Lu, W., Cowan, A.J., Morscher, R.J., Lee, G., Liu, W., Tesz, G.J., Birnbaum, M.J., Rabinowitz, J.D., 2018. The Small Intestine Converts Dietary Fructose into Glucose and Organic Acids. Cell Metabolism 27, 351–361.e3. 10.1016/j.cmet.2017.12.016

Jang, C., Wada, S., Yang, S., Gosis, B., Zeng, X., Zhang, Z., Shen, Y., Lee, G., Arany, Z., Rabinowitz, J.D., 2020. The small intestine shields the liver from fructose-induced steatosis. Nat Metab 2, 586–593. 10.1038/s42255-020-0222-9

Kim, M., Astapova, I.I., Flier, S.N., Hannou, S.A., Doridot, L., Sargsyan, A., Kou, H.H., Fowler, A.J., Liang, G., Herman, M.A., 2017. Intestinal, but not hepatic, ChREBP is required for fructose tolerance. JCI Insight 2. 10.1172/jci.insight.96703

Kleinert, M., Clemmensen, C., Hofmann, S.M., Moore, M.C., Renner, S., Woods, S.C., Huypens, P., Beckers, J., de Angelis, M.H., Schürmann, A., Bakhti, M., Klingenspor, M., Heiman, M., Cherrington, A.D., Ristow, M., Lickert, H., Wolf, E., Havel, P.J., Müller, T.D., Tschöp, M.H., 2018. Animal models of obesity and diabetes mellitus. Nat Rev Endocrinol 14, 140–162. 10.1038/nrendo.2017.161

Kokki, K., Lamichane, N., Nieminen, A.I., Ruhanen, H., Morikka, J., Robciuc, M., Rovenko, B.M., Havula, E., Käkelä, R., Hietakangas, V., 2021. Metabolic gene regulation by Drosophila GATA transcription factor Grain. PLoS Genet 17, e1009855. 10.1371/journal.pgen.1009855

Li, M.V., Chang, B., Imamura, M., Poungvarin, N., Chan, L., 2006. Glucose-dependent transcriptional regulation by an evolutionarily conserved glucose-sensing module. Diabetes 55, 1179–1189. 10.2337/db05-0822

Matsuda, H., Yamada, T., Yoshida, M., Nishimura, T., 2015. Flies without Trehalose *. Journal of Biological Chemistry 290, 1244–1255. 10.1074/jbc.M114.619411

Mattila, J., Havula, E., Suominen, E., Teesalu, M., Surakka, I., Hynynen, R., Kilpinen, H., Väänänen, J., Hovatta, I., Käkelä, R., Ripatti, S., Sandmann, T., Hietakangas, V., 2015. Mondo-Mlx Mediates Organismal Sugar Sensing through the Gli-Similar Transcription Factor Sugarbabe. Cell Reports 13, 350–364. 10.1016/j.celrep.2015.08.081

Öckerman, P.A., Lundborg, H., 1965. Conversion of fructose to glucose by human jejunum absence of galactose-to-glucose conversion. Biochimica et Biophysica Acta (BBA) - Enzymology and Biological Oxidation 105, 34–42. 10.1016/S0926-6593(65)80173-4

Overend, G., Luo, Y., Henderson, L., Douglas, A.E., Davies, S.A., Dow, J.A.T., 2016. Molecular mechanism and functional significance of acid generation in the Drosophila midgut. Sci Rep 6, 27242. 10.1038/srep27242

Pearl, L.H., Prodromou, C., 2006. Structure and mechanism of the Hsp90 molecular chaperone machinery. Annu Rev Biochem 75, 271–294. 10.1146/annurev.biochem.75.103004.142738

Phillips, M.D., Thomas, C.M., 2006. Brush border spectrin is required for early endosome recycling in Drosophila. J Cell Sci 119, 1361–1370. 10.1242/jcs.02839

Piper, M.D.W., Soultoukis, G.A., Blanc, E., Mesaros, A., Herbert, S.L., Juricic, P., He, X., Atanassov, I., Salmonowicz, H., Yang, M., Simpson, S.J., Ribeiro, C., Partridge, L., 2017. Matching Dietary Amino Acid Balance to the In Silico-Translated Exome Optimizes Growth and Reproduction without Cost to Lifespan. Cell Metabolism 25, 610–621. 10.1016/j.cmet.2017.02.005

Pustovaya, K., Venediktov, A., Soldatov, V., Kuzmin, E., Pokidova, K., Gartzeva, V., Payushina, O., Tsytsarev, V., Meglinski, I., Piavchenko, G., 2026. Recent insights into HSP70: proteostasis and beyond. Front. Mol. Biosci. 13. 10.3389/fmolb.2026.1791536

San Roman, A.K., Shivdasani, R.A., 2011. Boundaries, Junctions and Transitions in the Gastrointestinal Tract. Exp Cell Res 317, 2711–2718. 10.1016/j.yexcr.2011.07.011

Schottelius, A.J.G., Baldwin Jr., A.S., 1999. A role for transcription factor NF-kB in intestinal inflammation. Int J Colorect Dis 14, 18–28. 10.1007/s003840050178

Taylor, S.R., Ramsamooj, S., Liang, R.J., Katti, A., Pozovskiy, R., Vasan, N., Hwang, S.-K., Nahiyaan, N., Francoeur, N.J., Schatoff, E.M., Johnson, J.L., Shah, M.A., Dannenberg, A.J., Sebra, R.P., Dow, L.E., Cantley, L.C., Rhee, K.Y., Goncalves, M.D., 2021. Dietary fructose improves intestinal cell survival and nutrient absorption. Nature 597, 263–267. 10.1038/s41586-021-03827-2

van den Berg, L., Kokki, K., Wowro, S.J., Petricek, K.M., Deniz, O., Stegmann, C.A., Robciuc, M., Teesalu, M., Melvin, R.G., Nieminen, A.I., Schupp, M., Hietakangas, V., 2023. Sugar-responsive inhibition of Myc-dependent ribosome biogenesis by Clockwork orange. Cell Reports 42, 112739. 10.1016/j.celrep.2023.112739

Wang, X., Park, J., Susztak, K., Zhang, N.R., Li, M., 2019. Bulk tissue cell type deconvolution with multi-subject single-cell expression reference. Nat Commun 10,380. 10.1038/s41467-018-08023-x

Wu, M., Xu, L.-G., Zhai, Z., Shu, H.-B., 2003. SINK Is a p65-interacting Negative Regulator of NF-κB-dependent Transcription *. Journal of Biological Chemistry 278, 27072–27079. 10.1074/jbc.M209814200

Zhang, D., Tong, X., VanDommelen, K., Gupta, N., Stamper, K., Brady, G.F., Meng, Z., Lin, J., Rui, L., Omary, M.B., Yin, L., 2017. Lipogenic transcription factor ChREBP mediates fructose-induced metabolic adaptations to prevent hepatotoxicity. J Clin Invest 127, 2855–2867. 10.1172/JCI89934

Zhao, S., Jang, C., Liu, J., Uehara, K., Gilbert, M., Izzo, L., Zeng, X., Trefely, S., Fernandez, S., Carrer, A., Miller, K.D., Schug, Z.T., Snyder, N.W., Gade, T.P., Titchenell, P.M., Rabinowitz, J.D., Wellen, K.E., 2020. Dietary fructose feeds hepatic lipogenesis via microbiota-derived acetate. Nature 579, 586–591. 10.1038/s41586-020-2101-7

Zwick, R.K., Kasparek, P., Palikuqi, B., Viragova, S., Weichselbaum, L., McGinnis, C.S., McKinley, K.L., Rathnayake, A., Vaka, D., Nguyen, V., Trentesaux, C., Reyes, E., Gupta, A.R., Gartner, Z.J., Locksley, R.M., Gardner, J.M., Itzkovitz, S., Boffelli, D., Klein, O.D., 2024. Epithelial zonation along the mouse and human small intestine defines five discrete metabolic domains. Nat Cell Biol 26, 250–262. 10.1038/s41556-023-01337-z

